# Geographically widespread and novel hemotropic mycoplasmas and bartonellae in Mexican free-tailed bats and sympatric North American bat species

**DOI:** 10.1101/2024.02.08.577874

**Authors:** Daniel J. Becker, Kristin E. Dyer, Lauren R. Lock, Beckett L. Olbrys, Shawn A. Pladas, Anushka A. Sukhadia, Bret Demory, Juliana Maria Nunes Batista, Micaela Pineda, Nancy B. Simmons, Amanda M. Adams, Winifred F. Frick, M. Teague O’Mara, Dmitriy V. Volokhov

**Affiliations:** School of Biological Sciences, University of Oklahoma, Norman, OK, USA; Department of Biological Sciences, Southeastern Louisiana University, Hammond, LA, USA; Department of Preventive Veterinary Medicine and Animal Health, School of Veterinary Medicine and Animal Science, University of São Paulo, São Paulo, Brazil; Department of Mammalogy, Division of Vertebrate Zoology, American Museum of Natural History, New York, NY, USA; Bat Conservation International, Austin, TX, USA; Department of Ecology and Evolutionary Biology, University of California Santa Cruz, Santa Cruz, CA, USA; Department of Migration, Max Planck Institute of Animal Behavior, Radolfzell, Germany; Smithsonian Tropical Research Institute, Panama City, Panama; Center for Biologics Evaluation and Research, U.S. Food and Drug Administration, Silver Spring, MD, USA

**Keywords:** hemoplasmas, *Tadarida brasiliensis*, migration, One Health

## Abstract

Bacterial pathogens remain poorly characterized in bats, especially in North America. We describe novel (and in some cases panmictic) hemoplasmas (10.5% positivity) and bartonellae (25.5% positivity) across three colonies of Mexican free-tailed bats (*Tadarida brasiliensis*), a partially migratory species that can seasonally travel hundreds of kilometers. Molecular analyses identified three novel *Candidatus* hemoplasma species most similar to another novel *Candidatus* species in Neotropical molossid bats. We also detected novel hemoplasmas in sympatric cave myotis (*Myotis velifer*) and pallid bats (*Antrozous pallidus*), with sequences in the latter 96.5% related to *C*. Mycoplasma haemohominis. We identified nine *Bartonella* genogroups, including those in cave myotis with 96.7% similarity to *C*. Bartonella mayotimonensis. We also detected *Bartonella rochalimae* in migratory Mexican free-tailed bats, representing the first report of this human pathogen in the Chiroptera. The seasonality and diversity of these bacteria observed here suggest that additional longitudinal, genomic, and immunological studies in bats are warranted.

## Introduction

Bats have been intensively sampled for viral pathogens, with species in this mammalian order hosting multiple viruses with high virulence in humans (1). However, bats remain understudied for bacterial pathogens, which can be significant for their impacts on both human health and bat morbidity and even mortality (2, 3). Hemotropic mycoplasmas (hemoplasmas) and bartonellae are facultative intracellular bacteria of special interest in bats, given their high prevalence and substantial genetic diversity (4, 5). For example, sampling of Neotropical bat communities has identified many common and co-circulating genotypes of these bacteria (6–8). Surveys in Oceania and Europe have also supported plausible zoonotic transmission of these bacteria from bats to humans, including *Candidatus* Mycoplasma haemohominis and *C*. Bartonella mayotimonensis (9, 10). Greater characterization of these bacteria across global bat diversity (over 1,470 species) is therefore warranted to inform infection risks for both bats and humans, although little surveillance has thus far been conducted in North American bats (11, 12).

Although flight enables high mobility of bats, relatively few bat species undertake long-distance migrations (e.g., between maternity and wintering grounds) (13). In North America, Mexican free-tailed bats (*Tadarida brasiliensis*) display highly variable migratory strategies (14). The southwestern United States contains both non-migratory and migratory populations, with some individuals traveling hundreds to over 1,000 kilometers between wintering grounds in Mexico to northern maternity colonies in Oklahoma, Kansas, and Colorado (14–16). Other colonies across the species range include year-round residents (17, 18). This variation in migratory behavior could shape patterns of infection, including the seasonal dispersal of bacterial pathogens across landscapes to naïve hosts. Hemoplasmas have not yet been detected in this bat species (4, 19), and bartonellae have only been minimally described in the southernmost part of the bat species’ geographic range (i.e., Chile and Argentina) (20).

Here, we conducted an initial characterization of hemoplasmas and bartonellae in Mexican free-tailed bats across multiple populations and seasons. Our goals were to identify novel pathogens in this bat species and to test for differences in prevalence among colonies that differ in migratory strategy and across the bat annual cycle. We also tested whether pathogen lineages were unique to each geography or if migration may facilitate panmixia. Lastly, we used this opportunity to perform a pilot characterization of these pathogens in sympatric bat species.

## Material and Methods

### Wild bat sampling

We sampled three North American colonies of Mexican free-tailed bats in 2021 and 2022 to compare infections among migratory strategies and provide an initial assessment of pathogen seasonality. We sampled non-migratory individuals in southeastern Louisiana (17), focusing on a colony in Pine Grove of approximately 1,000 bats, in the non-breeding season (October 2021, *n*=5) and maternity season (July 2022, *n*=10). We also sampled the partially migratory population of Bracken Cave near San Antonio, Texas, which hosts tens of millions of this bat species in the maternity season and declines to approximately 10,000 bats in winter (16, 21). We sampled the maternity season (August 2021, *n*=20) and mid-to-late winter (December 2021 and March 2022; *n*=9 and *n*=19). We also sampled a fully migratory colony at the Selman Bat Cave near Freedom, Oklahoma, where this maternity roost holds up to 100,000 bats during summer and is empty in winter (16, 22). We sampled bats monthly, from April to September 2022, spanning spring arrival, the maternity season, and fall migration (*n=*146). In the same Oklahoma site, we also sampled four cave myotis (*Myotis velifer*), one hoary bat (*Lasiurus cinereus*), one Townsend’s big-eared bat (*Corynorhinus townsendii*), and two pallid bats (*Antrozous pallidus*).

Bats were captured with hand nets and mist nets while emerging from or returning to roosts and placed in individual cloth bags. Bats were identified to species by morphology and identified by sex, reproductive status, and age (23). Blood (<1% body mass) was sampled by lancing the propatagial vein using 27G and 30G needles and collected with heparinized capillaries. Blood was preserved on Whatman FTA cards and held at room temperature until −20°C storage at the University of Oklahoma (OU). Sampling was approved by the Institutional Animal Care and Use Committees of OU (2022-0198) and Southeastern Louisiana University (0064), with permits from the Texas Parks and Wildlife Department (SPR-0521-063), Louisiana Department of Wildlife and Fisheries (WDP-21-101), and Oklahoma Department of Wildlife Conservation (ODWC, 10567389). All bats were released after sampling at the capture site.

### Molecular diagnostics

We extracted genomic DNA from blood using QIAamp DNA Investigator Kits (Qiagen). To determine hemoplasma presence, we used PCR targeting the partial 16S and 23S rRNA genes (Table S1; (6, 7, 24, 25), with amplicons purified and sequenced at Psomagen. For DNA samples positive for the 16S or 23S rRNA genes, we also attempted to amplify the partial *rpoB* gene, using primers newly designed for this study (Table S1). To determine the presence of bartonellae, we used nested PCR targeting the partial *gltA* gene (Table S1; (26)), with amplicons purified with Zymo kits (DNA Clean & Concentrator-5, Zymoclean Gel DNA Recovery) and sequenced at the North Carolina State University Genomic Sciences Laboratory. We included blank FTA card punches and ultrapure water as extraction and negative controls, respectively, in all PCRs. Hemoplasma PCRs used *Candidatus* Mycoplasma haemozalophi as a positive control, but we did not include positive controls for *Bartonella* spp. to reduce cross-contamination risks from nested PCR; instead, amplicons of expected size (∼300 bp) were identified during gel electrophoresis. Sequences are available on GenBank through accessions OQ407831–50, OR783320–23, and PQ465198 (*Mycoplasma* spp. 16S rRNA); OQ359160–75 (*Mycoplasma* spp. 23S rRNA); OQ554332–38 (*Mycoplasma* spp. *rpoB*); and PP317862–72 (*Bartonella* spp. *gltA*).

### Statistical analysis

We analyzed infection states using generalized linear models (GLMs) or generalized additive models (GAMs) with binary response in R. All GLMs were fit using mean bias reduction methods with the *brglm2* package (27), whereas GAMs were fit using restricted maximum likelihood and the *mgcv* package (28). For each of our two pathogens, we fit the following four GLMs. The first model was fit to all data and compared the odds of infection among Louisiana, Texas, and Oklahoma bats. The second model was limited to the early-to-mid non-breeding season and compared odds of infection between Texas and Louisiana bats. The third model was limited to the late overwintering period and spring arrival to compare odds of infection between Texas and that subset of Oklahoma bats. The fourth model was limited to Texas and included reproductive status (only females were reproductive) and month to test demographic effects and seasonality (i.e., maternity season through late overwintering). Lastly, for our GAMs, we fit a similar model for each pathogen to the Oklahoma data, with a cyclic cubic smooth of month to assess a different aspect of infection seasonality (i.e., spring arrival until onset of fall migration).

### Phylogenetic analysis

We used NCBI BLASTn to identify related mycoplasma (16S rRNA, 23S rRNA, *rpoB*) and bartonellae sequences (*gltA*), which we aligned with our sequences and reference sequences using MUSCLE. We used MrBayes for phylogenetic analysis, with each gene tree run for 20,000,000 generations using a GTR+I+G model. BLASTn was implemented in Geneious, whereas MUSCLE and MrBayes were implemented using NGPhylogeny.fr (29). We delineated genotypes of hemoplasmas and genogroups of bartonellae based on pairwise similarity among sequences and clustering on their phylogenies, using established criteria for defining novel bacterial lineages (6, 30). For hemoplasmas, we also used multi-loci data to propose novel *Candidatus* species when the same genotype was identified in at least two samples using 16S rRNA and one other marker (i.e., 16S rRNA and 23S rRNA, 16S rRNA and *rpoB*) (7, 31).

We conducted two tests to assess if pathogen lineages were unique to each Mexican free-tailed bat colony, which would suggest geographically constrained transmission dynamics. First, we used chi-squared tests with *p* values generated by a Monte Carlo procedure to quantify associations between geography and pathogen lineage assignments. Next, for any lineages identified in multiple colonies with sufficient sample size, we derived matrices of spatial and phylogenetic distance among sequenced PCR-positive samples and used Mantel tests with the *vegan* R package to assess isolation by distance (32). These tests used 1,000 randomizations.

## Results

### Migratory and seasonal effects on bat bacterial infection

We detected hemoplasmas in 21 of 209 Mexican free-tailed bats when targeting the partial 16S or 23S rRNA genes (10.5%, 95% CI: 7.1−15.4%). Sequencing of the 23S rRNA gene showed three other bats (two from Texas and one from Oklahoma) had non-hemotropic mycoplasmas (OQ359169–70, OQ359174) most related to *Mycoplasma muris* (97.5% sequence identity). We detected bartonellae in 53 of 208 tested bats (25.5%, 95% CI: 20–31.8%). Only six Mexican free-tailed bats were coinfected by bartonellae and any mycoplasmas (2.9%, 95% CI: 1.3–6.1%). Hemoplasmas were detected in all three Mexican free-tailed bat colonies, while bartonellae were only detected in the Texas and Oklahoma colonies. PCR positivity data are fully available in the Pathogen Harmonized Observatory (PHAROS): https://pharos.viralemergence.org/ (33).

Across all Mexican free-tailed bats (model 1), the odds of infection differed by colony for bartonellae (*χ*^*2*^ = 9.72, *p <* 0.01) but not hemoplasmas (*χ*^*2*^ = 0.02, *p =* 0.99; Figure 1). When comparing only the resident and partially migratory populations in the non-breeding season (model 2), Louisiana and Texas bats did not differ in the odds of either infection (hemoplasmas: *χ*^*2*^ = 1.21, *p =* 0.27; bartonellae: *χ*^*2*^ = 0, *p =* 1). When comparing only the partially and fully migratory populations in spring (model 3), we did not detect colony differences for hemoplasmas (*χ*^*2*^ = 0, *p =* 1) or bartonellae (*χ*^*2*^ = 0.11, *p =* 0.74). When assessing risk factors of infection in the partially migratory Texas colony (model 4), we found no evidence of seasonal or demographic effects for either pathogen (Table S2). However, when assessing these predictors for the fully migratory Oklahoma colony across the full 2022 occupancy period, we identified significant seasonality in infection (Table S3). Prevalence increased from spring arrival and peaked in the maternity season (i.e., June 2022) for both bacteria, declining into fall migration (Figure 1).

**Figure 1.**
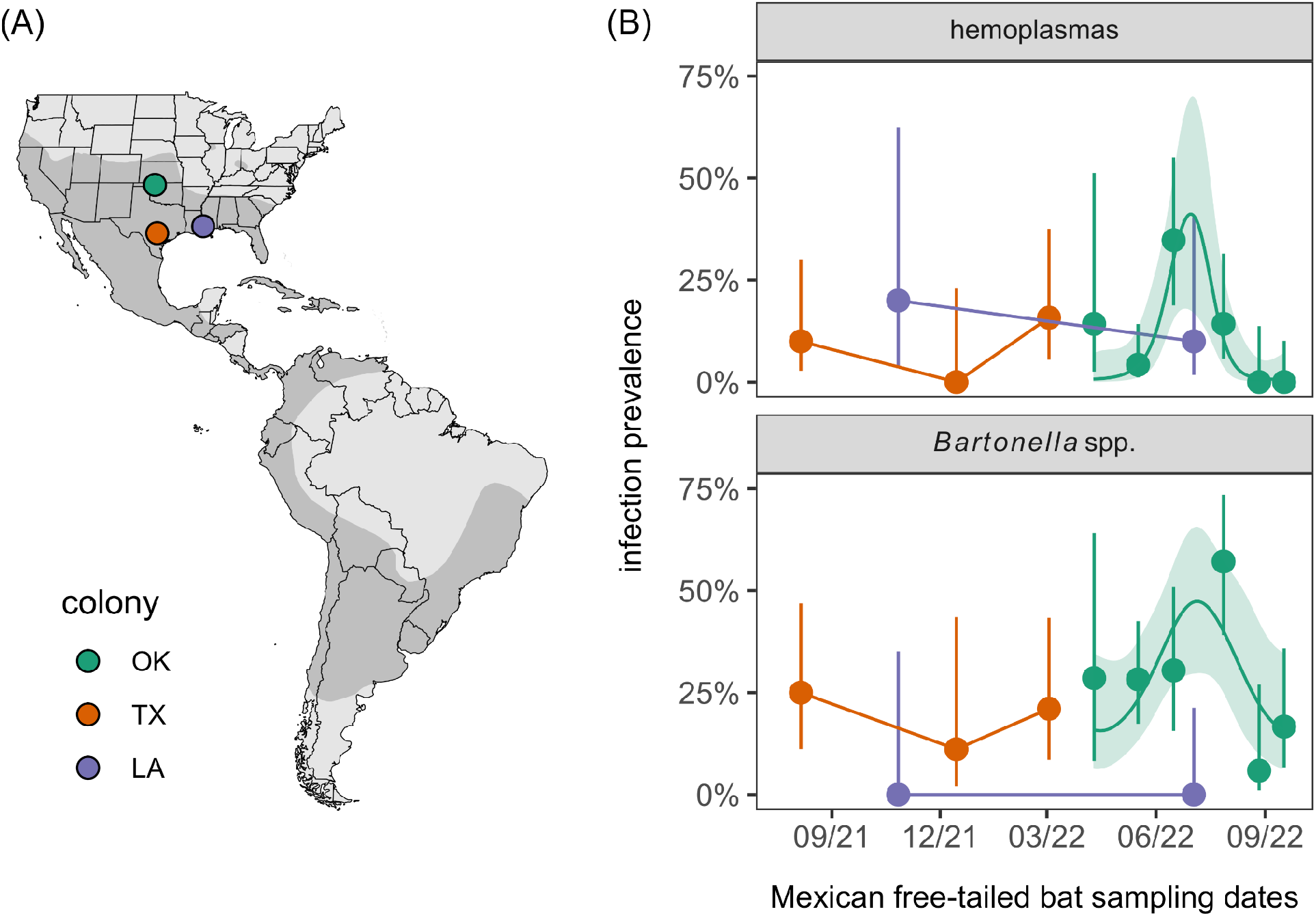
(A) Sampled Mexican free-tailed bat (*Tadarida brasiliensis*) colonies relative to the host distribution in the Americas. (B) Hemoplasma and *Bartonella* spp. infection prevalence across months and colonies; segments denote 95% confidence intervals using Wilson’s method. For Oklahoma bats,

### Genetic diversity of Mexican free-tailed bat hemoplasmas

Sequencing of 16S rRNA amplicons revealed four hemoplasma genotypes specific to Mexican free-tailed bats (i.e., TB1–4; Figure 2, Table 1). The TB1 genotype was 97% similar to the MR1 genotype that we earlier isolated from another molossid bat (*Molossus nigricans*) in Belize (e.g., MH245174) (6), and TB1 was found in all three sampled populations. In contrast, TB2 was only detected in the Texas colony and was ∼96–97% similar to hemoplasmas from carnivores and rodents and to the Belize bat MR1 genotype (6, 34, 35). Both TB3 and TB4 were only detected in Oklahoma and were ∼99% similar to MR1. We also detected the PPM1 genotype, originally found in *Pteronotus mesoamericanus* and *P. fulvus* in Belize (6, 7), in the Oklahoma colony.

**Table 1.**
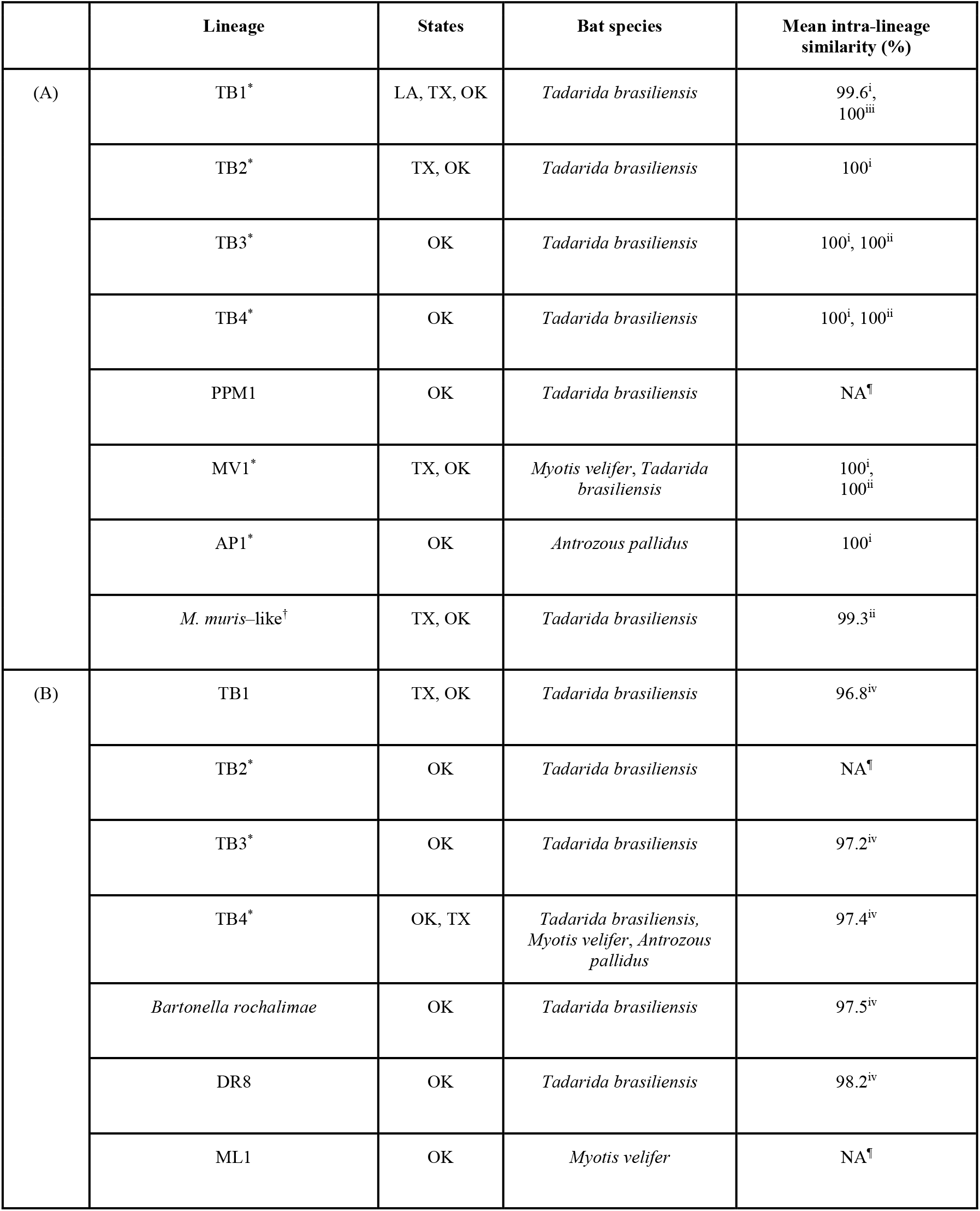
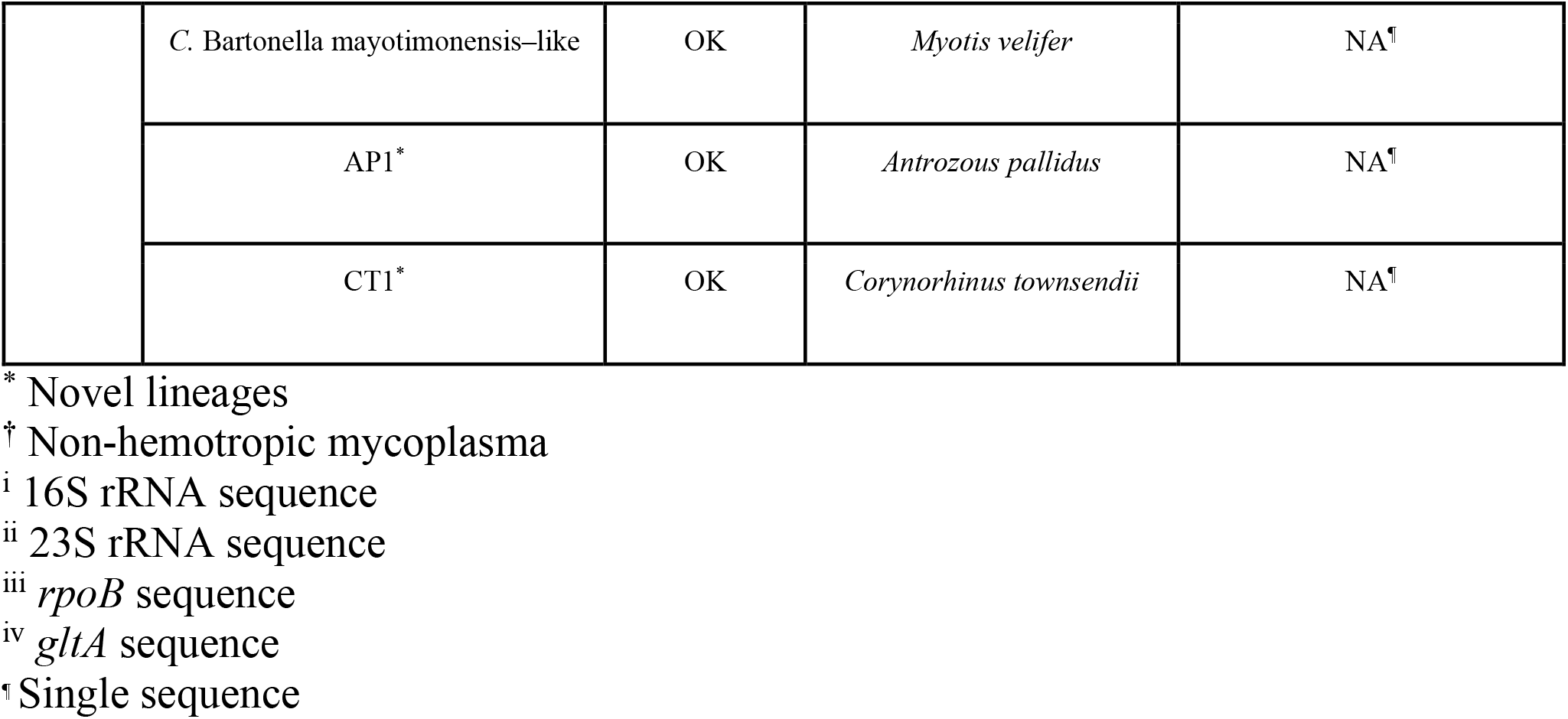
*Mycoplasma* spp. (A) and *Bartonella* spp. (B) lineages identified from Louisiana, Texas, and Oklahoma bats during this study (2021–2022). Lineages are given with their host species, locations, and mean intra-genotype (mycoplasmas) or intra-genogroup (bartonellae) sequence similarity from the partial 16S rRNA, 23S rRNA, *rpoB*, or *gltA* gene sequences identified here.

**Figure 2.**
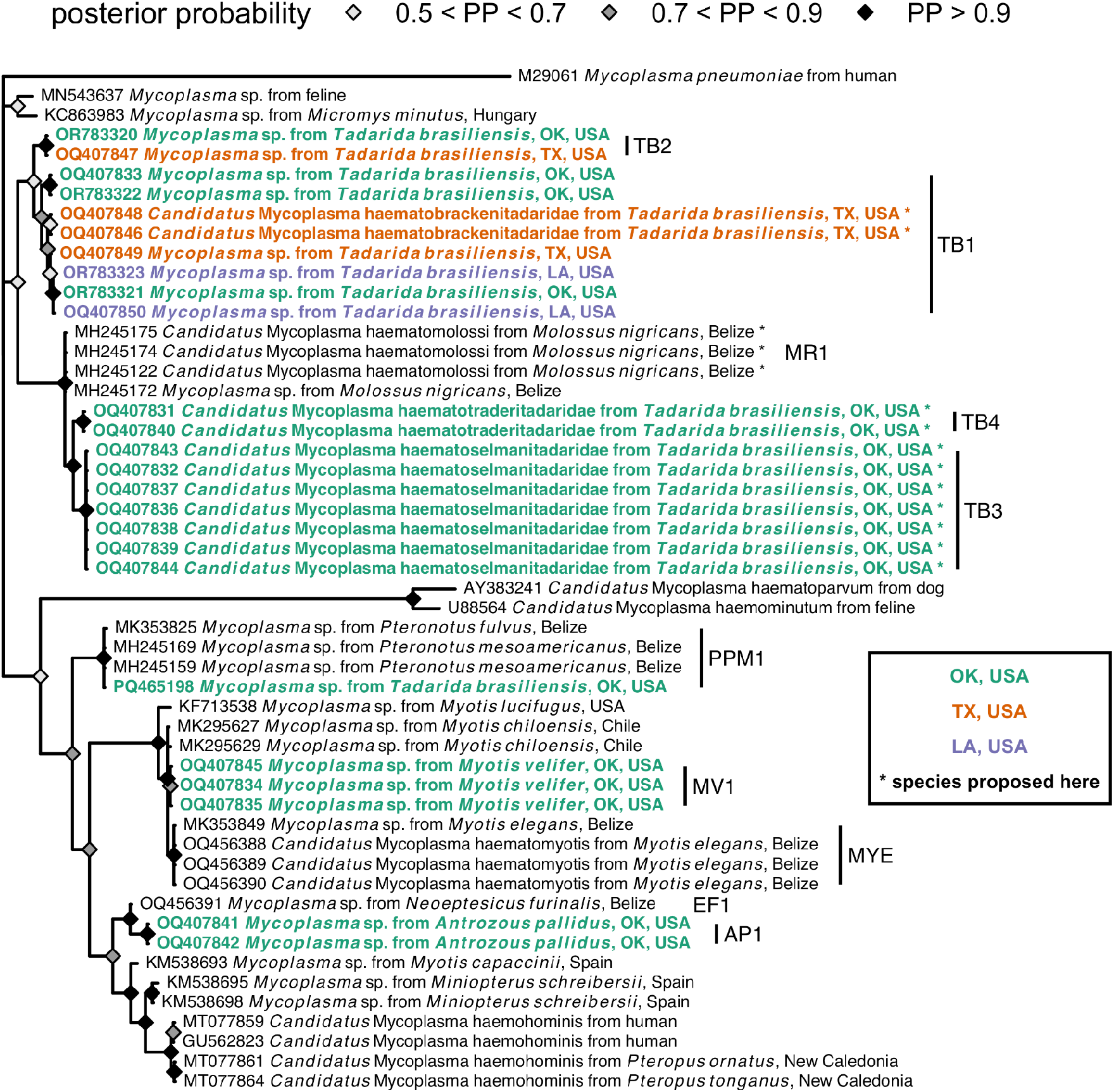
Consensus Bayesian phylogeny of the partial 16S rRNA hemoplasma sequences from this study (highlighted in bold and colored by geography; see Table 1 for genotype assignments) and reference sequences from bats and other mammals. Nodes are colored by posterior probability (nodes with less than 50% support are not shown). Hemoplasmas with *Candidatus* species names proposed here are indicated by asterisks and have paired 23S rRNA or *rpoB* sequences (see Figures S1 and S3).

Amplification of the 23S rRNA gene from two bats (one each from Texas and Oklahoma) also found a genotype we initially detected in cave myotis (i.e., MV1; Figure S1) and the above *Mycoplasma muris*–like (non-hemotropic) genotype. These seven mycoplasma genotypes were not associated with geography (*χ*^*2*^ = 13.07 *p* = 0.38; Figure S2). When considering the one genotype observed in multiple colonies and in more than one bat per site (TB1), we also found little support for isolation by distance with the 16S rRNA phylogeny (Mantel *r* = 0.35, *p* = 0.10).

Amplification of paired partial 23S rRNA (Figure S1) and/or *rpoB* (Figure S3) genes for samples belonging to these 16S rRNA genotypes suggested at least three novel *Candidatus* hemoplasma species circulate in Mexican free-tailed bats. Based on 100% identity of two *rpoB* sequences (OQ554335–36) and 100% identity of paired 16S rRNA sequences included in the TB1 genotype, first detected in Bracken Cave (OQ407846, OQ407848), we propose the name *C*. Mycoplasma haematobrackenitadaridae sp. nov. Similarly, given 99.98% identity among seven 23S rRNA sequences (OQ359161–65, OQ359168, OQ359172) and 100% identity in paired 16S rRNA sequences included in the TB3 genotype from the Selman Bat Cave (OQ407832, OQ407836–39, OQ407843–44), we propose the name *C*. M. haematoselmanitadaridae sp. nov. Lastly, given 100% identity of two 23S rRNA sequences (OQ359160, OQ359166) and 99.8% identity in paired 16S rRNA sequences from the TB4 genotype (also identified from the Selman Bat Cave; OQ407831, OQ407840), we propose the name *C*. M. haematotraderitadaridae sp. nov. (Figures 2 and S1), based on the stream running adjacent to the bat cave (Traders Creek) (36).

Given the similarity of Mexican free-tailed bat hemoplasma 16S rRNA sequences to those from molossid bats sampled in Belize (6), we also attempted to amplify the 23S rRNA and *rpoB* genes from *Molossus nigricans* sampled in 2017 and 2018 in Belize that previously tested positive for the MR1 and MR2 genotypes (6). We re-extracted DNA from four FTA cards and applied the same additional PCR protocols described earlier. We obtained partial 23S rRNA and *rpoB* sequences for two (OQ518943–44) and three (OQ554329–31) *M. nigricans*, respectively. Based on 100% inter-sequence similarity of the *rpoB* sequences and high 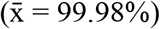 identity of paired 16S rRNA sequences (MH245122, MH245172, MH245174), we propose the name *C*. Mycoplasma haematomolossi sp. nov. to designate this novel hemoplasma (Figures 2 and S3).

### Genetic diversity of Mexican free-tailed bat bartonellae

Sequencing of *gltA* amplicons next revealed at least six *Bartonella* genogroups circulating in Mexican free-tailed bats (Figure 3, Table 1). The first genogroup was detected in both Texas and Oklahoma, with sequences from all three sampled months across 2021 and 2022 from Bracken Cave but only from May and June in the Selman Bat Cave. The *gltA* sequences were ≥99.6% similar to those recently detected in Mexican free-tailed bats in Argentina (KX986617) (20), such that we consider these sequences to all form the TB1 genogroup. The TB2–4 genogroups were only found in the Oklahoma colony. TB2 was identical to sequences from streblid bat flies in Mexico (e.g., ≥99.7% identity to MF988072) (37), while TB3 represents a novel genogroup with ∼92% similarity to *Bartonella vinsonii* (e.g., MK984790) (38). Likewise, TB4 was novel and distantly related (∼91%) to *Bartonella quintana* (i.e., Z70014) (39). Oklahoma bats also harbored g*ltA* sequences with 97–100% identity to *Bartonella rochalimae* in fleas from foxes (OQ436435) (40); these *B. rochalimae* sequences were only detected in summer months. Lastly, Oklahoma bats had *gltA* sequences in the same clade as bartonellae originally found in vampire bats (*Desmodus rotundus*; DR8 genogroup) (26). These six *Bartonella* genogroups were not associated with geography (*χ*^*2*^ = 8.20, *p* = 0.16; Figure S4), and we also found no support for isolation by distance for the TB1 genotype with our *glt*A phylogeny (Mantel *r* = -0.04, *p* = 0.59).

**Figure 3.**
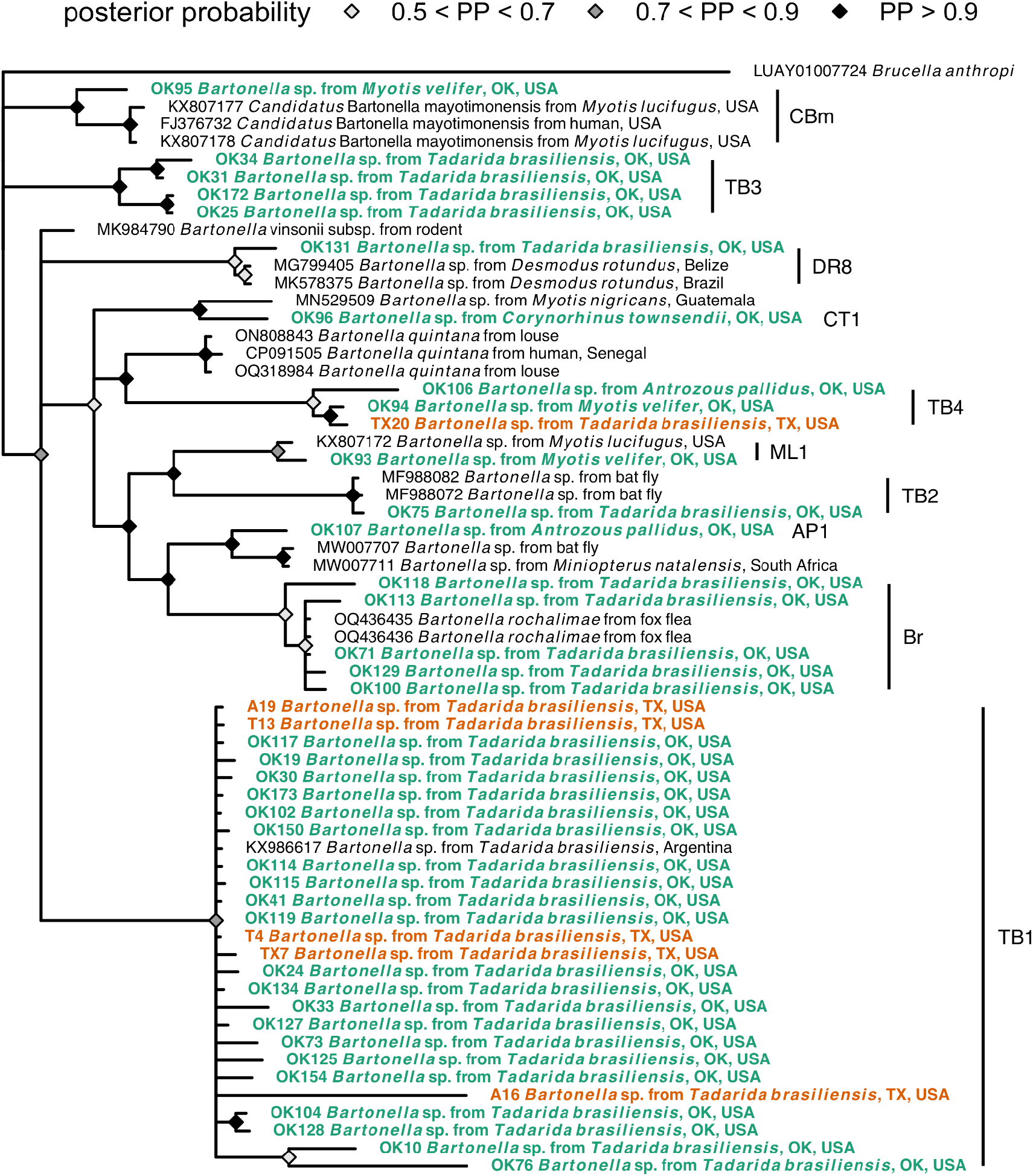
Consensus Bayesian phylogeny of the partial *gltA Bartonella* spp. sequences from this study (highlighted in bold and colored by geography; see Table 1 for genogroup assignments) and reference sequences from bats, other mammals, and ectoparasites. Nodes are colored by posterior probability (nodes with less than 50% support are not shown).

### Bacterial infections of sympatric bat species

Opportunistic sampling of other bats in Oklahoma revealed further bacterial diversity (Table 1). Three of the four cave myotis and both pallid bats tested positive for hemoplasmas, whereas three of three cave myotis, both pallid bats, and the single Townsend’s big-eared bat tested positive for bartonellae; the single hoary bat tested negative for both bacterial pathogens.

For hemoplasmas (Figure 2), we identified a single novel hemoplasma genotype in each PCR-positive species (i.e., MV1 and AP1), with 16S rRNA sequences most closely related (i.e., ≥98% similarity) to previously detected genotypes in vesper bats from Chile and Belize (e.g., EF1 and MYE) (6, 19). Notably, 16S rRNA sequences of the AP1 genotype were 96.5% similar to those of *Candidatus* Mycoplasma haemohominis (i.e., GU562823), whereas those of the MV1 genotype were only ∼94% similar. 23S rRNA and *rpoB* sequences from cave myotis (OQ359173, OQ554337) were entirely novel (<85% similarity to GenBank sequences), with the former 100% similar to select 23S rRNA sequences identified from our Mexican free-tailed bats.

For bartonellae (Figure 3), all three positive cave myotis had their own genogroup. One *gltA* sequence was ∼98% similar to that first identified in little brown bats (*Myotis lucifugus*) elsewhere in North America (i.e., KX807172), here denoted the ML1 genogroup. Another cave myotis had the TB4 genotype. The final cave myotis *gltA* sequence clustered within a clade of *Candidatus* Bartonella mayotimonensis sequences (∼96%), isolated from a human endocarditis patient in Iowa, USA (FJ376732) and other little brown bats (KX807177–9) (11, 41). For pallid bats, one sequence also belonged to the TB4 genogroup, whereas the other formed the novel AP1 genogroup most related (∼95%) to bartonellae from Natal long-fingered bats (*Miniopterus natalensis*) and their bat flies in South Africa (e.g., MW007711 and MW007707) (42). Lastly, the single positive Townsend’s big-eared bat hosted a unique genogroup (CT1) only ∼94% similar to bartonellae found from *Myotis nigricans* in Guatemala (e.g., MN529509) (5).

## Discussion

Hemoplasmas and bartonellae are emerging as model systems for studying bacterial infections in bats (5, 7), but their infection dynamics and diversity remain poorly characterized, notably in North American systems (11, 12). We here demonstrate novel diversity of hemoplasmas and bartonellae in bats in the south-central United States, including the circulation of lineages of both pathogens with clear infection seasonality in a migratory colony of Mexican free-tailed bats. Such work provides the foundation for further empirical studies to elucidate the transmission dynamics of these bacteria, their pathogenicity in bats, and their possible zoonotic risk.

Our findings suggest relatively common infection with site-specific and panmictic bacterial lineages. Within both bacterial genera, most lineages found in Mexican free-tailed bats were restricted to a single site, indicating spatially constrained transmission. However, we also detected lineages in multiple sites, such as the TB1 hemoplasma genotype in Louisiana, Texas, and Oklahoma; TB1 and MV1 hemoplasma genotypes as well as the *Mycoplasma muris–*like genotype in Texas and Oklahoma; and TB1 *Bartonella* genogroup in Texas and Oklahoma (for which sequences were nearly identical to those from *Tadarida brasiliensis* in Argentina) (20).

Such results may be explained by migratory connectivity in Mexican free-tailed bats, for which regional migrations spanning hundreds to over 1,000 kilometers have been well-characterized in North America, including between the Selman Bat Cave and Bracken Cave (14, 15, 43); this suggests the migratory behavior of this species can enhance bacterial dispersal. However, given the presence of these bacterial lineages in migratory and non-migratory populations (i.e., Louisiana) and at the extremes of the bat range (e.g., over 5,000 km between the Selman Bat Cave and the Argentina site for the TB1 *Bartonella* genogroup), these results also suggest the ancestral spread of these bacteria and limited selection pressure on lineages across the bat range.

Future studies are needed to identify the migratory routes of Mexican free-tailed bats, especially for understanding the origins of possibly zoonotic bacterial lineages and the potential for these bats to disperse infection during spring and fall migrations. Researchers could capitalize on advances in tracking small vertebrates for long periods, such as use of absorbent sutures, to ensure lightweight radiotags stay attached to bats for the duration of migration and winter (44).

Such work is also needed to assess if these infections negatively impact bat migration trajectory and success, as observed for blood pathogens in migratory songbirds (45). Longitudinal studies would also inform such analyses, as our data from one full occupancy season in the Oklahoma colony suggest bacterial prevalence peaks in the maternity season. Additional seasonal sampling is needed to assess how infection risk varies across the full migratory cycle, if prevalence tracks bat population size and/or ectoparasite intensity, and whether infections are sufficiently common in autumn to facilitate their dispersal with migration. Further genetic analyses could also inform patterns of bat connectivity and pathogen spread. For example, the TB2 *Bartonella* genogroup from Oklahoma Mexican free-tailed bats showed 100% identity to bartonellae from bat flies from Morelos, Hidalgo, and Jalisco in central Mexico (37), spanning the likely wintering sites of this bat species (14, 15). Similarly, previous analyses of *Trypanasoma cruzi* from this same Oklahoma bat population detected lineages similar to those along the Texas–Mexico border, further showing possible southern origins of infection and high pathogen dispersal capacity (46).

Additional molecular and immunological studies are also needed to better characterize these novel bat bacterial pathogens and their health impacts. We identified 16S rRNA and *gltA* sequences with moderate-to-high similarity to zoonotic pathogens such as *C*. Mycoplasma haemohominis, *C*. Bartonella mayotimonensis, and *Bartonella rochalimae* (9, 41, 47). For the former two pathogens, our bat sequences were ∼96% similar to zoonotic lineages, likely indicating divergence from a common ancestor at least tens of million of years ago (7, 48). *B. rochalimae* has been found in cats and dogs (49), and our detections in Mexican free-tailed bats indicate a broadening of host range into bats. Our sequences showed ∼97–100% similarity to those from fox fleas (40), relevant given likely flea transmission (49) and detection of fleas in the Oklahoma bat population where these sequences were found (Dyer, personal communication). Generation of whole genomes for our novel bat pathogens could inform their zoonotic risk, both by better linking them to cryptic human infections (9) and by facilitating machine learning models that predict zoonotic potential from genomic composition, as applied for viruses (50). Other -omics analyses could also elucidate whether these bacterial infections are pathogenic in bats themselves. In addition to assessing impacts of infection on migration outcomes as noted above, approaches such as transcriptomics and proteomics could test if bats have a pronounced immune response to these bacterial infections or appear largely tolerant (51). Such studies could be especially informative when comparing immunity between migratory and non-migratory periods, which could test whether long-distance migration may disrupt immune tolerance in bats.

Lastly, Mexican free-tailed bats and their sympatric bat species provide several important ecosystem services, including but not limited to predating on crop pests and contributing to the tourism economy from bat flight watching (52, 53). Understanding the prevalence, genetic diversity, and pathogenicity of bacterial pathogens in bats can inform One Health approaches that emphasize conservation measures to promote bat, domestic animal, and human health (54).

## Acknowledgements

This work was supported by the National Institute of General Medical Sciences of the National Institutes of Health (P20GM134973), Louisiana Board of Regents Support Fund, Louisiana Biomedical Research Network, and Research Corporation for Science Advancement (RCSA). This work was conducted via Subawards No. 28365 and 29018, part of a USDA Non-Assistance Cooperative Agreement with RCSA Federal Award No. 58-3022-0-005. Additional support was provided by the Edward Mallinckrodt, Jr. Foundation and University of Oklahoma (Data Institute for Societal Challenges, Vice President for Research and Partnerships). Bat Conservation International provided in-kind support. We thank William Caire, Melynda Hickman, and the ODWC for site access; the Selman Living Laboratory for fieldwork support; and Konstantin Chumakov for laboratory support. We thank one anonymous reviewer for helpful feedback.

