## Supplemental Material for "Geographically widespread and novel hemotropic mycoplasmas and bartonellae in Mexican free-tailed bats and sympatric North American bat species"

Table S1. PCR primers and amplification parameters used in this study
Table S2. GLM results for Mexican free-tailed bats from Bracken Cave, Texas
Table S3. GAM results for Mexican free-tailed bats from from Selman Bat Cave, Oklahoma
Figure S1. Consensus Bayesian phylogeny of *Mycoplasma* 23S rRNA sequences
Figure S2. Cross-tabulation of *Mycoplasma* genotypes by geography
Figure S3. Consensus Bayesian phylogeny of *Mycoplasma rpoB* sequences
Figure S4. Cross-tabulation of *Bartonella* genotypes by geography

Table S1. PCR primers and amplification parameters used in this study.

| Primers | Sequence | Organism and gene | Expected product (bp) | Temperature (ºC) / time (seconds) | | | | Cycles | Ref |
|---|---|---|---|---|---|---|---|---|---|
| | | | | Initial denaturation | Denaturation | Annealing | Extension | | |
| CS 443f | GCTATGTCTGCATTCTATCA | Bartonellae *gltA* | ~700 | 95/120 | 95/30 | 48/30 | 72/120 | 40 | 1 |
| CS 1210r | GATCYTCAATCATTTCTTTCCA | | | | | | | | |
| Bhcs 781p | GGGGACCAGCTCATGGTGG | | ~300 | 95/180 | 95/30 | 55/30 | 72/120 | 40 | 2 |
| Bhcs 1137n | AATGCAAAAAGAACAGTAAACA | | | | | | | | |
| UNI_16S_hemoFnew | TGAATAAGTGACAGCWAACTATGTGCC | Hemoplasma 16S rRNA | ~850–900 | 95/300 | 95/50 | 60/60 | 72/60 | 55 | 3* This study |
| UNI_16S_hemoR | GACGGGCGGTGTGTACAAGACCTG | | | | | | | | |
| UNI_rpoB_hemoF1 | CCTAAYTTRARYATWMGKGACGTTCACTATTC | Hemoplasma *rpoB* (primer set 1)** | ~785 | 95/300 | 95/50 | 55/60 | 72/60 | 55 | This study |
| UNI_rpoB_hemoR1_1 | GAAGAMARRATAATDGCATCYTCATAGTTGTA | | | | | | | | |
| UNI_rpoB_hemoF1 | CCTAAYTTRARYATWMGKGACGTTCACTATTC | Hemoplasma *rpoB* (primer set 2)** | ~1280 | 95/300 | 95/50 | 55/60 | 72/90 | 55 | This study |
| UNI_rpoB_hemoR1_2 | ACAGGAGTWCCATCYTCYARRTAWGGCAT | | | | | | | | |
| UNI_23S_Myc_Ur_cladeF | CCCAGACCATKGGGYAAGCCTA | Hemoplasma 23S rRNA | ~1500–80 | 95/300 | 95/50 | 58/60 | 72/90 | 55 | 3 |
| UNI_23S_Myc_Ur_cladeR | GAGACAGTCAAGAGATGGTTACAC | | | | | | | | |

*Primers were slightly modified based on available hemoplasma 16S rRNA gene data

**Both sets of primers were used to amplify the hemoplasma *rpoB* gene. The *rpoB* primers were designed based on the conserved sequences found within the *rpoB* gene sequences of known hemotropic mycoplasmas and closely related *Mycoplasma* species.

1. Birtles RJ, Raoult D. Comparison of partial citrate synthase gene (*gltA*) sequences for phylogenetic analysis of *Bartonella* species. *International Journal of Systematic and Evolutionary Microbiology*. 1996;46(4):891-7.

2. Norman AF, Regnery R, Jameson P, Greene C, Krause D. Differentiation of *Bartonella*-like isolates at the species level by PCR-restriction fragment length polymorphism in the citrate synthase gene. *Journal of Clinical Microbiology*. 1995;33(7):1797-803.

3. Volokhov DV, Norris T, Rios C, Davidson MK, Messick JB, Gulland FM, Chizhikov VE. Novel hemotrophic mycoplasma identified in naturally infected California sea lions (*Zalophus californianus*). *Veterinary Microbiology*. 2011;149(1-2):262-8.

Table S2. Results of GLMs with mean bias reduction for hemoplasma and *Bartonella* spp. positivity in Mexican free-tailed bat samples from Bracken Cave in Texas ($n$ = 48; model 4). Reference levels include bats sampled in August 2021 and non-reproductive bats.

| | hemoplasmas | | | *Bartonella* spp. | | |
|---|---|---|---|---|---|---|
| Parameter | OR | $\|z\|$ | $p$ | OR | $\|z\|$ | $p$ |
| Intercept | | 2.25 | 0.02 | | 1.96 | 0.05 |
| December 2021 | 0.40 | 0.51 | 0.61 | 0.74 | 0.25 | 0.80 |
| March 2022 | 1.63 | 0.45 | 0.65 | 1.22 | 0.22 | 0.83 |
| Reproductive | 1.53 | 0.33 | 0.74 | 2.67 | 0.95 | 0.34 |

Table S3. Results of GAMs for hemoplasma and *Bartonella* spp. positivity in Mexican free-tailed bat samples from Selman Bat Cave in Oklahoma ($n$ = 146 and $n$ = 145, respectively). Non-reproductive bats serve as the reference (only females were reproductive). Predictors are presented with model coefficients or estimated degrees of freedom (EDF) and test statistics

| | hemoplasmas | | | | | *Bartonella* spp. | | | | |
|---|---|---|---|---|---|---|---|---|---|---|
| Term | OR | $\|z\|$ | EDF | $\chi^2$ | $p$ | OR | $\|z\|$ | EDF | $\chi^2$ | $p$ |
| Intercept | | 4.67 | | | <0.01 | | 2.77 | | | 0.01 |
| Reproductive | 0.39 | 1.36 | | | 0.17 | 0.79 | 0.53 | | | 0.60 |
| s(Week) | | | 2.10 | 14.02 | 0.001 | | | 1.75 | 6.68 | 0.02 |

Figure S1. Consensus Bayesian phylogeny of partial 23S rRNA mycoplasma sequences from this study (highlighted in bold and colored by geography; see Table 1 for genotype assignments) and reference sequences from bats and other mammals. Nodes are colored by posterior probability (nodes with less than 50% support are not shown). Hemoplasmas with *Candidatus* species names proposed here are indicated by asterisks and have paired 16S rRNA sequences in Figure 1.

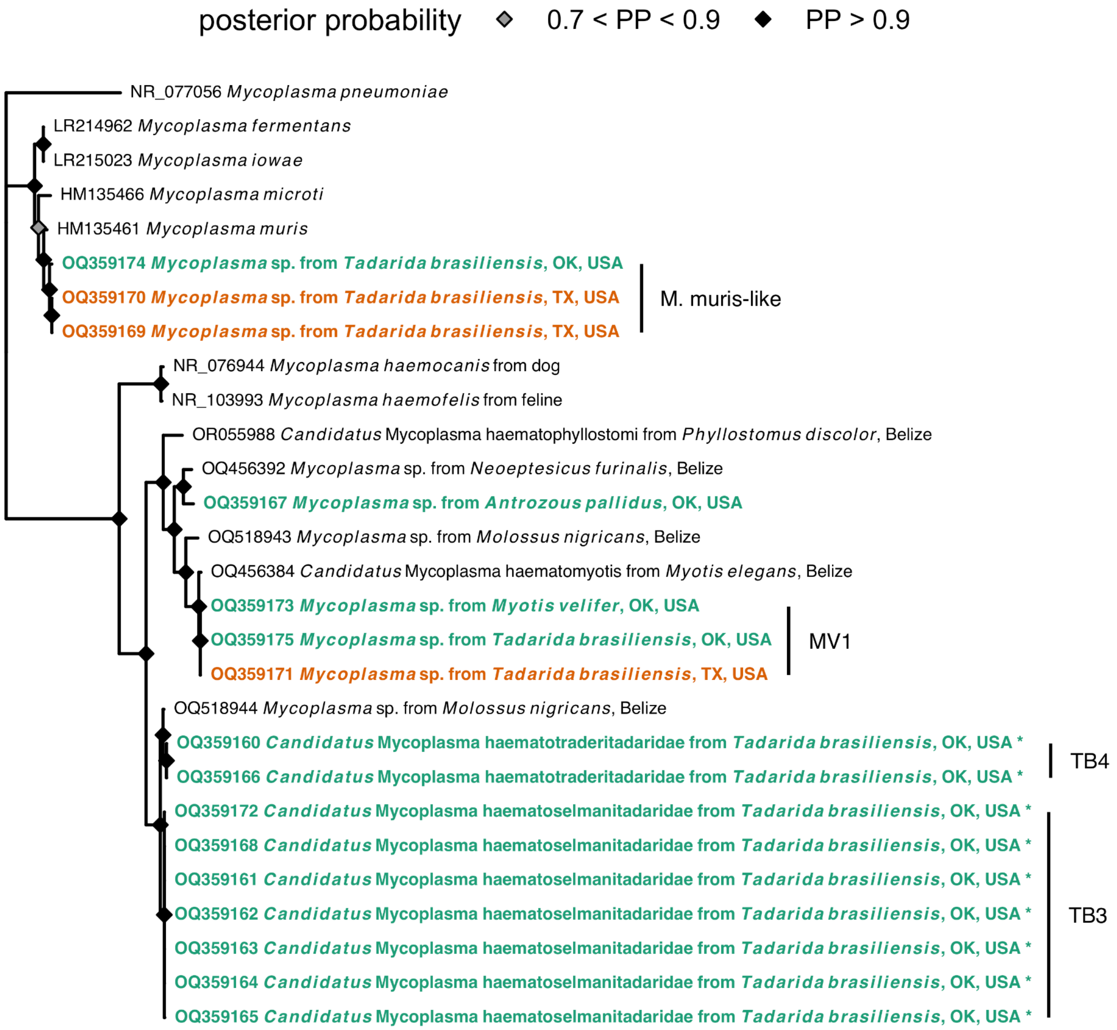

Figure S2. Cross-tabulation of *Mycoplasma* genotypes by geography, including all bat species.

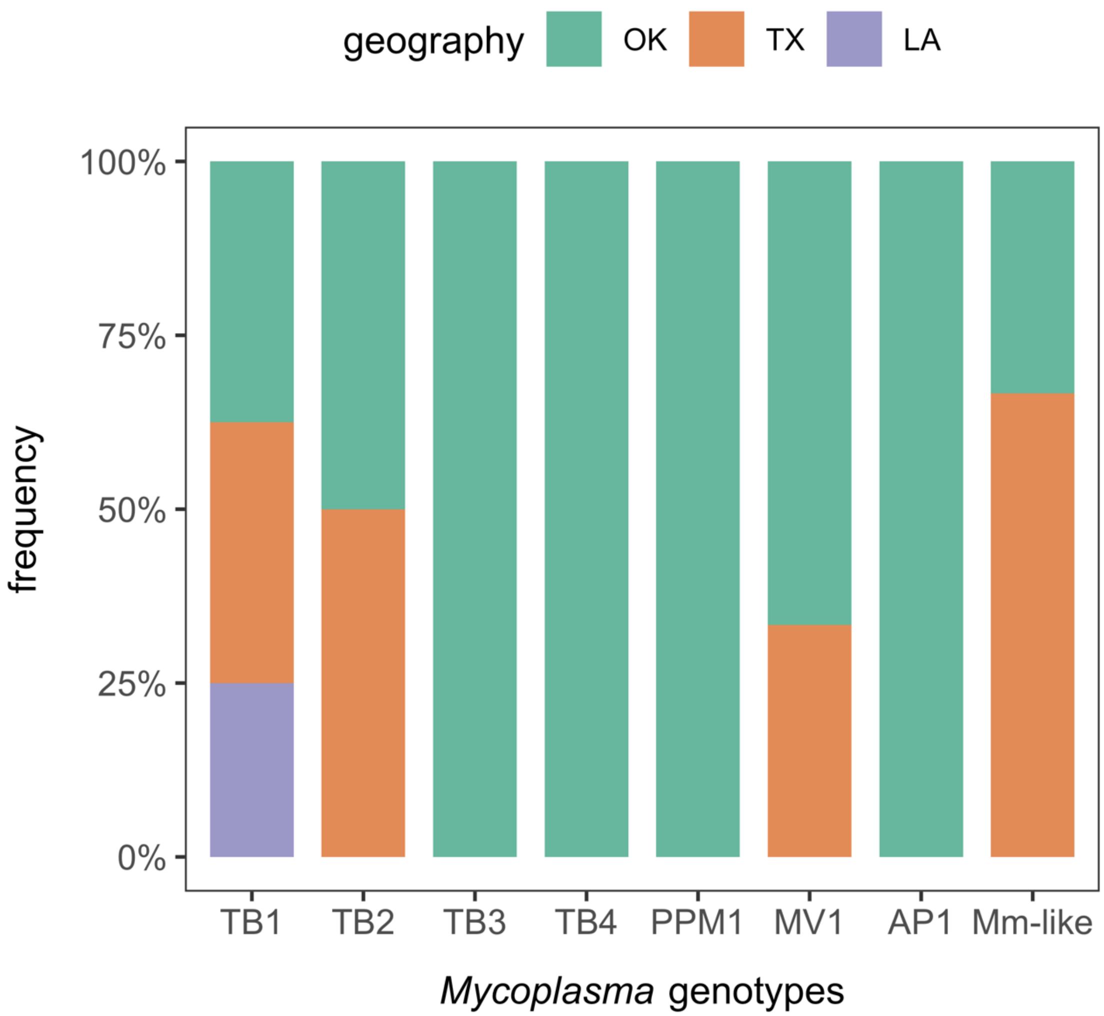

Figure S3. Consensus Bayesian phylogeny of partial *rpoB* mycoplasma sequences from this study (highlighted in bold and colored by geography; see Table 1 for genotype assignments) and reference sequences from bats and other mammals. Nodes are colored by posterior probability (nodes with less than 50% support are not shown). Hemoplasmas with *Candidatus* species names proposed here are indicated by asterisks and have paired 16S rRNA sequences in Figure 1.

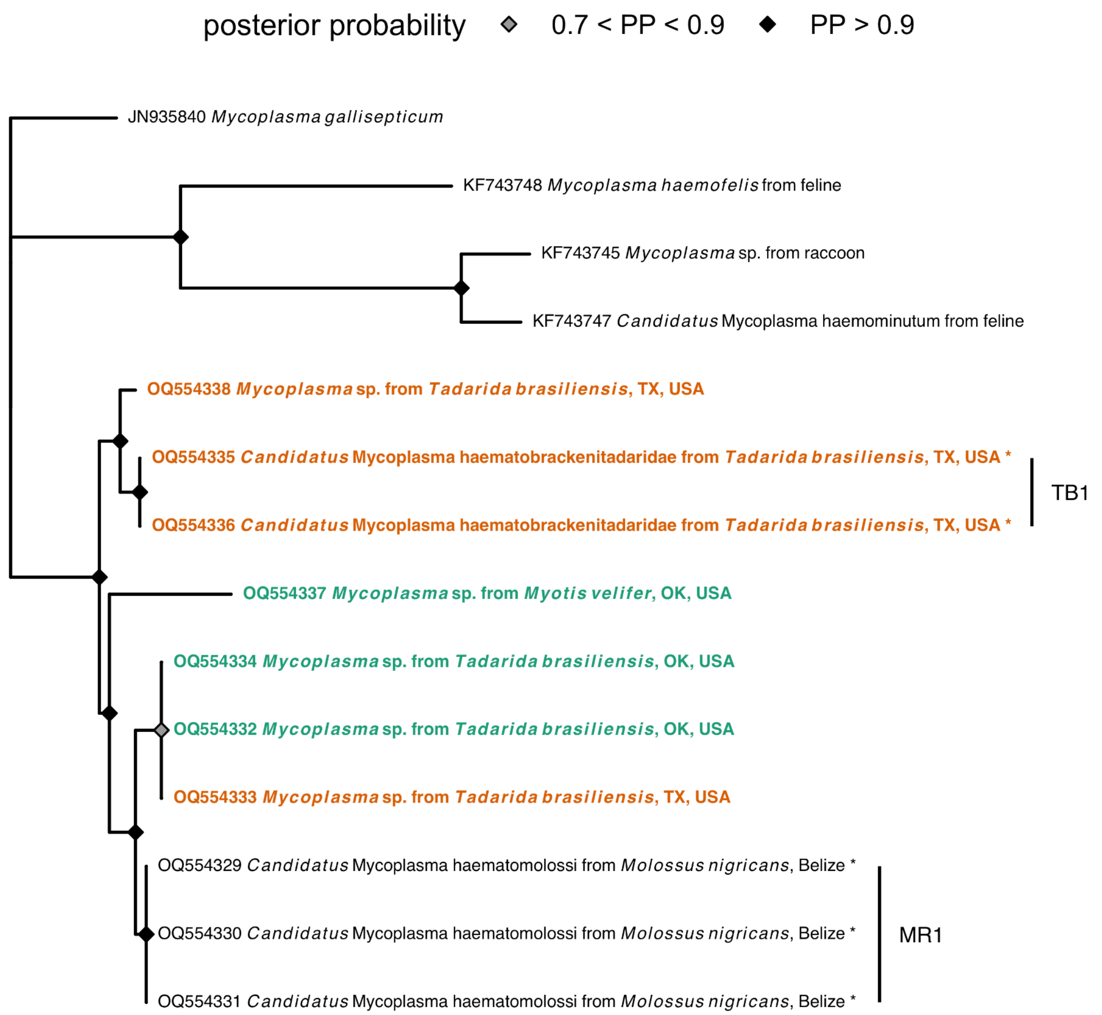

Figure S4. Cross-tabulation of *Bartonella* genotypes by geography, including all bat species.

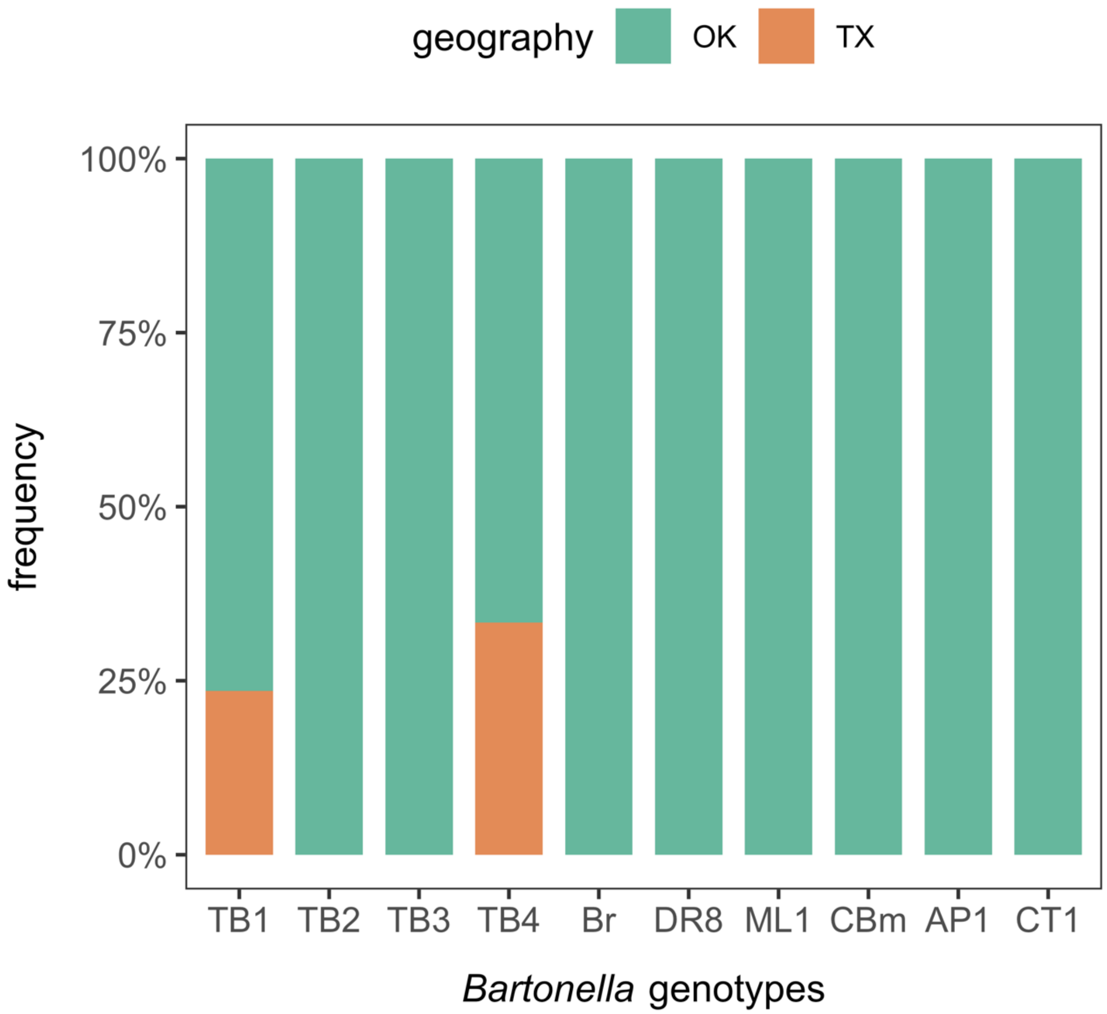
